# Identification of Y-chromosome turnover in newts fails to support a sex chromosome origin for the *Triturus* balanced lethal system

**DOI:** 10.1101/2024.11.04.621952

**Authors:** James France, Wiesław Babik, Milena Cvijanović, Katarzyna Dudek, Ana Ivanović, Tijana Vučić, Ben Wielstra

## Abstract

Non-recombining regions of the genome often have profound effects on evolution, resulting in phenomena such as sex chromosomes and supergenes. Amongst the strangest examples are balanced lethal systems, such as that found in newts of the genus *Triturus*. These systems halve reproductive output, and the evolution of such a deleterious trait is difficult to explain. For *Triturus* an intriguing model proposes that the balanced lethal system evolved from an ancestral Y-chromosome. To test this hypothesis, we identify the Y-chromosome of *Triturus* and verify whether it, or the balanced lethal system, is homologous to the Y-chromosome of its sister genus *Lissotriton*, which does not possess the balanced lethal system. We identify a set of candidate Y-linked markers in *T. ivanbureschi* and place them on a high-density linkage map that we construct with 7,233 RADseq markers. We validate male specificity of the markers across the genus, and then place both the *Triturus* and *Lissotriton* Y-linked regions within previously constructed target capture linkage maps that include genes linked to the balanced lethal system. We observe that neither the *Triturus* balanced lethal system, nor the *Triturus* Y-chromosome are homologous to the *Lissotriton* Y-chromosome. This is the first molecular evidence of a transition between Y-chromosome systems within salamanders. However, unless additional sex chromosome turnover events are involved, our data does not support a sex chromosome origin of the balanced lethal system.

## Introduction

In crested and marbled newts (the genus *Triturus*) the first and largest chromosome occurs in two distinct versions (termed 1A and 1B), which do not undergo recombination along most of their length (Callan et al., 1960). All adult *Triturus* newts possess one copy of each of these versions (genotype 1A/1B). However, as each offspring receives one of each chromosome pair randomly from both of its parents, half of the offspring will inherit two copies of the same version of chromosome 1 (1A/1A or 1B/1B). These offspring fail to develop normally and die before hatching (Macgregor & Horner, 1980). The lethality of a balanced lethal system derives from the presence of unique non-functional alleles of essential genes within the non-recombining region of each version of the chromosome (Muller, 1918). In *Triturus* our previous research has identified two private sets of genes – one present on each of chromosomes 1A and 1B (France et al., 2024a; de Visser et al., 2024a). Any embryo that inherits a 1A/1A or 1B/1B genotype will completely lack functional alleles for one set of these genes, whereas embryos with the 1A/1B genotype will possess at least one functioning allele for all genes. As each version of the chromosome is required to compensate for the deficiencies of the other, neither can be selected against and both are maintained at equal frequencies.

Several other cases of balanced lethal systems have been described in widely divergent lineages, including central American *Drosophila* and plants of the genera *Isotoma* and *Oenothera* (Dobzhansky & Pavlovsky, 1955; James et al., 1990; Steiner, 1956). The repeated evolution of such a maladaptive trait, which results in the loss of 50% of reproductive output, is difficult to explain. However, there are several proposals which link the origin of balanced lethal system in *Triturus* newts to other phenomena characterized by suppressed recombination, such as supergenes (Berdan et al., 2022; Wielstra, 2020) and sex chromosomes (Wallace, 1984). Testing of these hypotheses may offer unique insight into how non-recombining regions of the genome can evolve into new roles systems which may exhibit with pronounced negative phenotypic effects.

A detailed model concerning the evolution of the *Triturus* balanced lethal system was developed by Grossen et al. (2012) (Fig. 1). Titled “A Ghost of Sex Chromosomes Past” it proposes that *Triturus*’ chromosome 1 evolved from an ancestral Y-chromosome. As sex chromosomes typically do not undergo recombination, outside of any pseudoautosomal regions, the ancestral chromosome would be free to split into two distinct lineages, Y_A_ and Y_B_, which would eventually become chromosomes 1A and 1B (Charlesworth et al., 2005). After the Y_A_ and Y_B_ lineages diverged, the model proposes that both began to accumulate lethal alleles. As long as these alleles were recessive they would not have been selected against, because their effect would have been masked by the presence of the X-chromosome. The model then enforces a climatic shift sufficient to override the masculinization factor present on the Y-chromosome. This results in some XY individuals having a female phenotype, and so creates the potential of offspring with a YY genotype. Any offspring with the genotypes Y_A_Y_A_ or Y_B_Y_B_ would be non-viable due to possessing two copies of one of the lethal alleles. However, if there were no shared lethal alleles present on both the Y_A_ and Y_B_ chromosomes, then individuals with the genotype Y_A_Y_B_ could survive. If the effect of temperature-induced sex reversal grew strong enough, all XY and even some YY individuals would develop as female. The resulting female-biased sex ratio would result in selection against the X-chromosome, eventually driving it extinct, leaving Y_A_Y_B_ as the only viable genotype, and so creating a balanced lethal system. Finally, the model proposes that the biased sex ratio leads to the evolution of a new masculinizing factor on an autosome, creating a new Y-chromosome.

**Figure 1:**
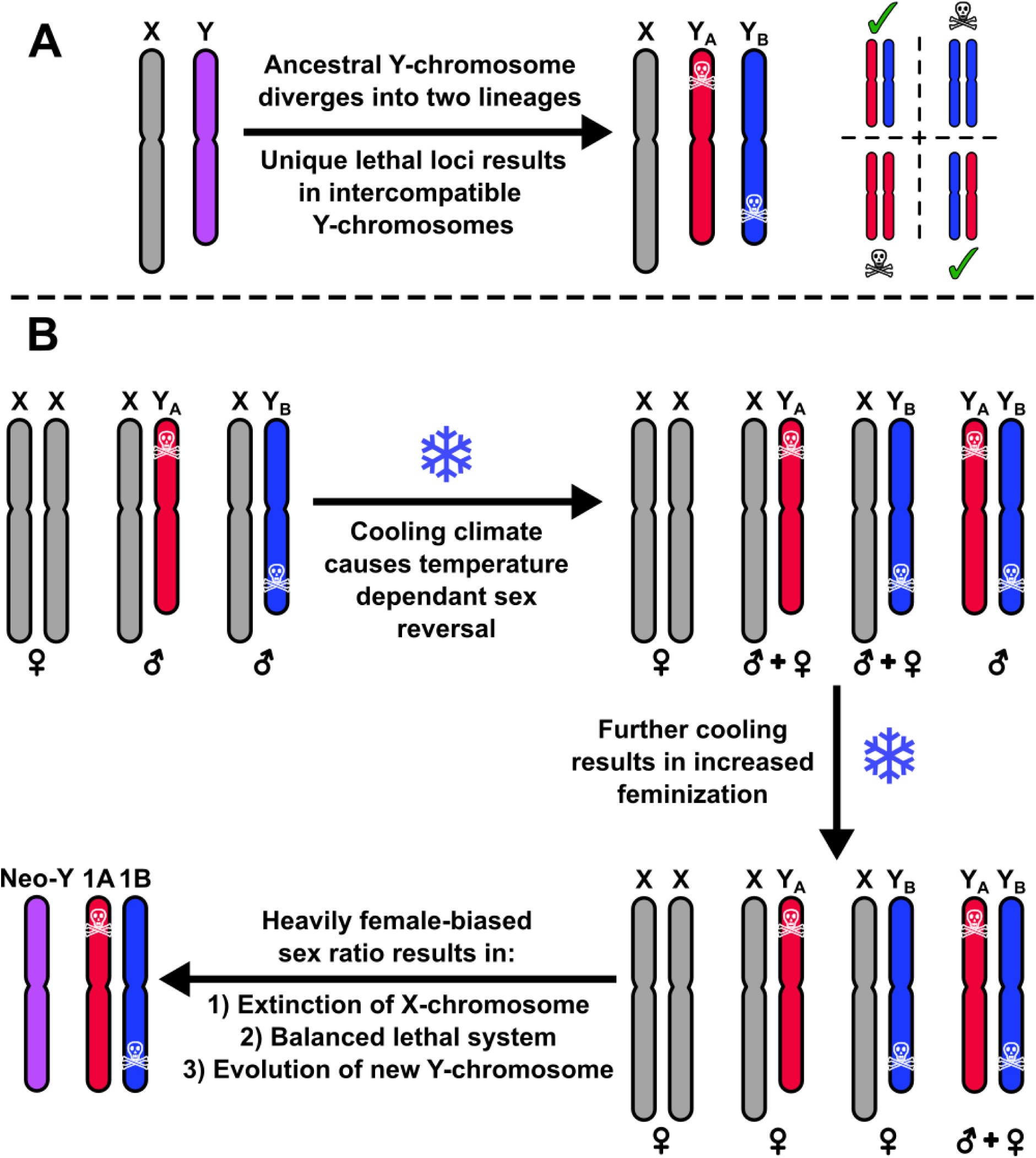
The hypothesised evolution of the *Triturus* balanced lethal system from a Y-chromosome (Grossen et al., 2012). A) The ancestral Y-chromosome diverges into two lineages, each other which possesses its own lethal alleles. Normally the Y-chromosomes cannot meet, but if they did only individuals with one copy of each of the two different lineages would be viable. B) As the climate cooled, temperature dependent sex reversal would result in XY females, producing the possibility of YY individuals. With drastic climate change even a YY genotype may not always be sufficient for masculinization, resulting in a sufficiently female biased sex-ratio that the X-chromosome is driven extinct, even though this results in a balanced lethal system.

While the evolution of a balanced lethal system via the mechanism proposed by Grossen et al. (2012) is prerequisite on a very particular coincidence of multiple specific factors, none of these are completely implausible in isolation. Y-chromosomes will tend to degenerate and accumulate lethal factors as a direct consequence of their lack of recombination (Charlesworth & Charlesworth, 2000). Temperature-induced sex reversal is common in many amphibians, including *Triturus* newts (Wallace & Wallace, 2000). The lethally homozygote but inter-compatible Y-chromosome lineages proposed are an almost exact analogue of the situation observed in guppies (Haskins et al., 1970). Furthermore, the evolution of new sex chromosomes is frequent in salamanders, as evidenced by the multiple transitions between XY and ZW systems within the order (Sessions, 2008).

A major virtue of the ‘’Ghosts of Sex chromosomes past” hypothesis is that it implies simple and readily testable predictions (Fig. 2). Firstly, because the proposed mechanism requires a sex chromosome turnover event, modern *Triturus* newts could not have retained the sex determination system that existed before the evolution of the balanced lethal system. Therefore, the Y-chromosome of *Triturus* should not be homologous to that of any relatives that diverged before the evolution of the balanced lethal system. Secondly, if *Triturus* chromosome 1 did previously function as a sex chromosome, then it should be homologous to the Y-chromosomes of other closely related newt genera - unless they had independently also lost the ancestral Y-chromosome.

**Figure 2:**
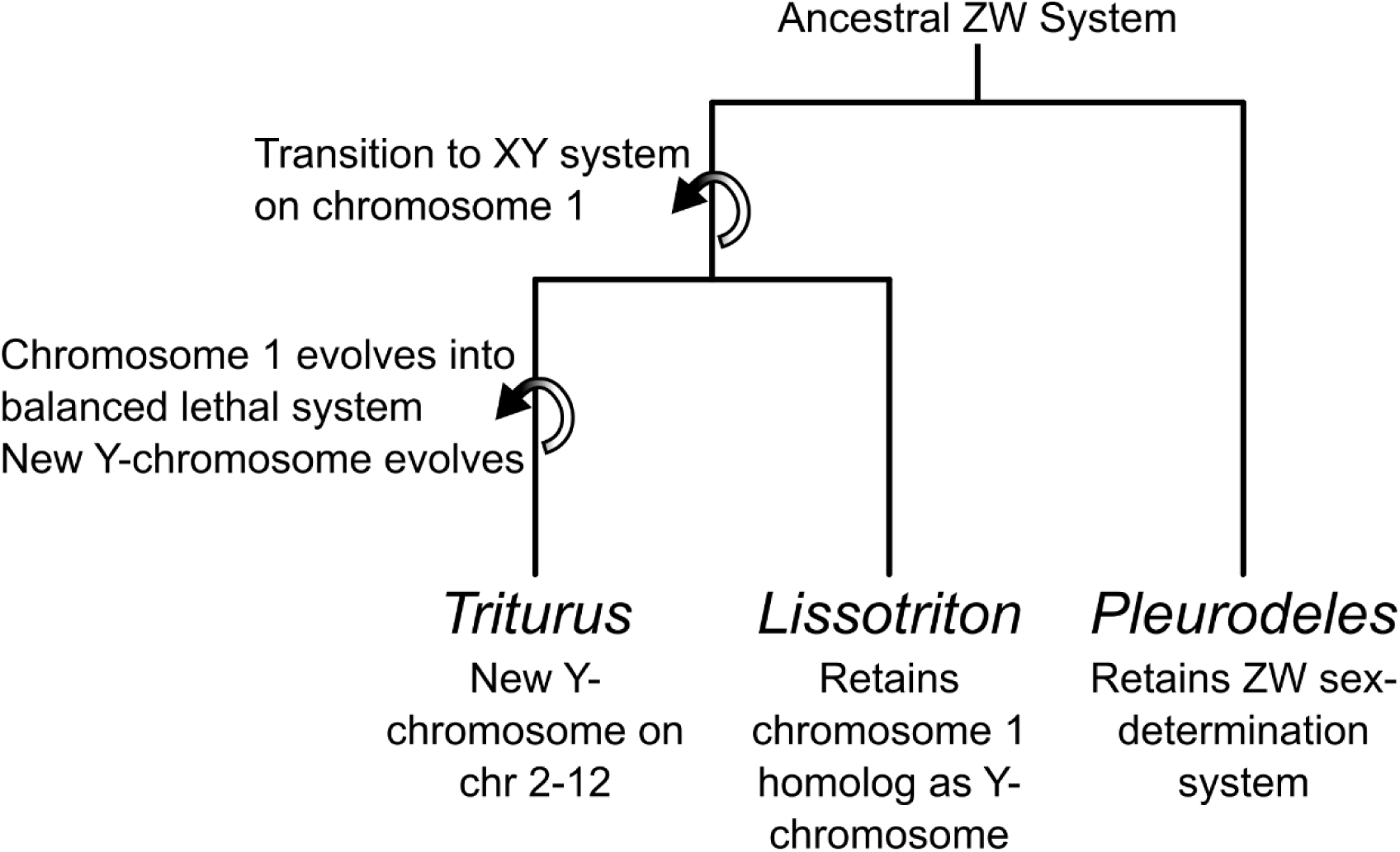
Schematic of sex chromosomes turnover events in newts required by the Y-chromosome origin hypothesis. A transition between an ancestral male-determining region on chromosome 1 to a new Y-chromosome is an essential feature of this mechanism. This implies that the Y-chromosome of *Lissotriton* cannot be homologous with that of *Triturus* and will instead likely be homologous with the *Triturus* balanced lethal system. Also indicated is transition from the ancestral salamander ZW system (which appears to be retained in the basal genus *Pleurodeles*) to the derived XY systems reported in *Triturus* and related newt genera such as *Lissotriton*.

In this study we test whether the Y-chromosome of *Triturus* is homologous to that of newts of its sister genus *Lissotriton* (Rancilhac et al., 2021), which does not possess the balanced lethal system. Cytological studies have identified largely homomorphic X and Y-chromosomes in both genera, but have not been able to determine if these, or any of the other 11 chromosome pairs, are homologous (Schmid et al., 1979; Sims et al., 1984). Molecular genomics is challenging due to the extremely large genome size of salamanders, estimated at approximately 30 Gbp in both *Triturus* and *Lissotriton* (Litvinchuk et al., 2007). We previously used RAD sequencing to identify Y-linked molecular markers for the smooth newt, *Lissotriton vulgaris* and place them within its genome (France et al., 2024b). Here, we apply the same strategy to the Balkan crested newt *Triturus ivanbureschi* to identify Y-linked molecular markers and position them within a RADseq based linkage map which we construct.

We then use two strategies to compare the Y-chromosomes of *Triturus* and *Lissotriton*. Firstly, we align the RADseq linkage maps for both genera with the genome assembly of the Iberian ribbed-newt *Pleurodeles waltl* (Brown et al., 2023) – we previously showed that the *Lissotriton vulgaris* Y-chromosome was homologous to chromosome 5 in *Pleurodeles waltl* (France et al., 2024b), if this is also the case for the *Triturus ivanbureschi* Y-chromosome it would indicate that the two genera both retain the sex chromosome of their common ancestor and so no sex chromosome turnover has occurred. Secondly, we utilize target capture based linkage maps that we previously constructed for both *Triturus* and *Lissotriton*, which show the position of genes linked to the balanced lethal system in *Triturus* and their homologs in *Lissotriton* (France et al., 2024a). We use PCR to screen the families used to construct these target capture linkage maps for the Y-linked RAD markers we identify in *Triturus* in this study (and previously in *Lissotriton*), allowing us to position the Y-linked RAD markers within the target capture maps. This allows us to identify any potential synteny between the Y-chromosomes of the two newt genera and the balanced lethal system.

## Methods & Materials

### Samples

For identification of candidate Y-linked markers, we performed RADseq on tissue samples of 60 adult, morphologically sexed *T. ivanbureschi* from Zli Dol (Pčinja district, Serbia). The RADseq linkage map was based on a family consisting of two adult T. *ivanbureschi* (one male, one female, also collected from Zli Dol) and 158 offspring (healthy and arrested embryos) of undetermined sex. The embryos were obtained in an experimental crossing under controlled laboratory conditions. For genus-wide validation of candidate markers via PCR we tested all other species within the genus recognized at time of sampling, including *T. anatolicus*, *T. carnifex, T. cristatus, T. dobrogicus, T. karelinii, T. macedonicus, T. marmoratus* and *T. pygmaeus* (Wielstra & Arntzen, 2016) – the recently recognised *T. rudolfi* (Arntzen, 2024) was not yet described at the time. For each of these species we used a single male-female pair, except for *T. carnifex*, where we used one pair each from both the Balkan and Italian lineages, which show high genetic divergence (Wielstra et al., 2021). A full list of samples used in this study is found in sup. Table S1.

### DNA extraction, library preparation and RAD-sequencing

Genomic DNA was extracted from the samples with the Promega Wizard™ Genomic DNA Purification Kit (Promega, Madison, WI, USA), according to the salt-based extraction protocol of Sambrook and Russel (2001). The Adapterama III High-Throughput 3RAD (Bayona-Vásquez et al., 2019) protocol was used to prepare RADseq libraries from 100 ng of sample DNA, using restriction enzymes *Eco*RI, *Xba*I, and *Nhe*I. Fragments in the range of 490-600 bp were excised using a Pippin Prep system (Sage Science, Beverly, MA, USA), the libraries were pooled equimolarly and 150 bp paired-end sequencing was performed by Novogene (Cambridge, UK) on the Illumina NovaSeq 6000 (Illumina Inc., San Diego, CA, USA) platform.

### RADseq data processing

The stacks package v2.54 (Catchen et al., 2013) was used for the processing of RADseq data obtained from the *T. ivanbureschi* samples. After demultiplexing and trimming via the process_radtags program the denovo_map.pl pipeline was used to assign reads to loci. For the linkage map family, the default settings were used without alteration. In particular, the parameter M, which determines the amount of divergence two reads may have while being assigned to the same putative locus, was kept at 2, maximising the number of loci recovered from the closely related sample set. For the adults of known-sex, the parameter M was set to 10, even though this reduces the number of loci recovered our experience with *Lissotriton* has shown this results in candidate Y-linked markers which are less likely to produce false positive results in female newts (France et al., 2024b).

### Developing candidate Y-linked from known-sex *T. ivanbureschi*

In order to select loci which were present only in male individuals, the BAM files produced by denovo_map.pl were used to create a matrix listing the coverage of all markers in each sample by employing the depth function of SAMtools (Li et al., 2009). This matrix was then filtered with a custom R script to produce a list of candidate Y-linked markers, present in at least 90% of male samples and absent in at least 90% of females. To reduce the chance of selecting markers with autosomal paralogs, which may result in false positive results in PCR assays, the candidate Y-linked markers were BLASTed (Camacho et al., 2009) against the catalogue of all markers produced by denovo_map.pl and paralogous hits with > 80% sequence similarity with a query coverage of > 25% recorded. Candidate markers were ranked based on absence of residual reads in females, number of potential paralogs and average read depth in males.

### Validation of candidate Y-linked markers via PCR

Primer 3 (Untergasser et al., 2012) was used to design primer pairs for the 12 highest ranked candidate markers, targeting an optimal primer length of 20 bp and melting temperature of 60°C. For each marker we attempted to design two sets of primers to amplify both a long (ca. 200 bp) and short (ca. 100 bp) fragment. The short fragment sequences were derived entirely from the forward reads of the RADseq data, whereas the long sequences bridged both forward and reverse reads. As the majority of read pairs were non-overlapping, the long fragments incorporate an additional sequence of unknown length.

The primers were initially tested by PCR in a male-female pair of *T. ivanbureschi*. The 2x QIAGEN multiplex master mix (QIAGEN B.V, Venlo, Netherlands) was used with a PCR protocol consisting of a 95°C hot start for 10 minutes, followed by 35 cycles of denaturation for 30 seconds at 95°C, 60 seconds annealing at 63°C and 45 seconds extension at 72°C, with a final extension of 10 minutes at 72°C. All primers were used at a final concentration of 0.1 µM.

Any primer pairs which showed amplification only in the male *T. ivanbureschi* were then tested in male-female pairs of *T. macedonicus* and *T. cristatus*. Primer pairs also showing male-specific amplification in these taxa were then tested in the remaining *Triturus* species. Finally, the best performing markers were selected, and a multiplex PCR designed, incorporating CDK-17 (de Visser et al., 2024b) which amplifies a product of 537 bp, as an autosomal control marker.

### RADseq linkage map construction and analysis

A joint VCF file produced by Stacks was filtered with VCFtools (Danecek et al., 2011) to exclude indels and SNPs with greater than 5% missing data, a mean depth of less than 10, or a minor allele frequency of less than 0.2. The thin function was then used to select a single SNP per locus. Separately, a custom R script was used to determine the coverage of the candidate Y-linked markers, identified in the known-sex adults, within each sample used for the linkage map. This data was then converted into presence/absence genotype calls with a custom R script, treating presence of the marker as an artificial SNP locus of genotype AT and absence as AA.

Lep-MAP 3 (Rastas, 2017) was then used to construct a linkage map. After the first stage of the pipeline (ParentCall2) the Y-linked presence/absence calls were appended to the output. Initial linkage groups were created with the SeparateChromosomes2 module, with a LOD limit of 20 (chosen as the number of linkage groups recovered rises rapidly with increasing LOD until 20 whereafter it plateaus) and distortion LOD set to 1. Unplaced markers were then added with the JoinSingles2All module with a LOD limit of 15. The markers were then ordered with the OrderMarkers2 module, using 12 merge iterations, 6 polish iterations, a minError value of 0.02, the scale setting M/N 2 and employing the sexAveraged option.

The sequences of the makers placed on the resulting linkage map were then blasted against the genome assembly of the Iberian ribbed newt (*Pleurodeles waltl*) (Brown et al., 2023), using a word size of 11 and requiring a minimum E value of 1e-20. Following the methodology of Purcell *et al*. (2014) results were then filtered to include only hits that exceeded the significance of the next highest ranked hit by at least five orders of magnitude. The hits that remained after filtering were visualised with an Oxford plot to show syntenic relationships between *Triturus* linkage groups and *P. waltl* chromosomes.

### Incorporation of Y-linked markers into target-capture linkage maps

In a previous study we used target capture to construct linkage maps based on ca. 7k coding genes for both *Triturus* (using an F_2_ *T. ivanbureschi* x *T. macedonicus* family) and *Lissotriton* (F_2_ *L. vulgaris* x *L. montandoni*), which allowed for the identification a set of genes associated with the balanced lethal system on *Triturus* chromosome 1, and their homologs in *Lissotriton* (France et al., 2024a). However, as none of the target capture markers were sex-linked, the location of the Y-linked regions could not be ascertained from these maps. To determine whether the Y-linked region of the *Lissotriton* genome was homologous to the balanced lethal system of *Triturus* (as would be expected if the balanced lethal system evolved from the shared ancestral sex chromosome), we needed to incorporate the Y-linked RADseq markers discovered in *L. vulgaris* (France et al., 2024b) into the target capture linkage map. To this end we used PCR to screen all samples from the family used to construct the *Lissotriton* target capture linkage map for the RAD marker lvY-51393-short, which is Y-linked and amplifies a product only in males. We also used this method to incorporate the *Triturus* Y-linked RAD marker TiY-384959-short, identified in this study, into the *Triturus* target capture linkage map. In both PCR screenings the primers for the Y-linked markers were multiplexed with those for CDK-17, to provide an autosomal control.

Following genotyping of offspring, presence/absence of the Y-linked marker was converted into pseudo-SNP genotyping calls in a manner similar to that described above, with samples that amplified the Y-linked band given an artificial SNP locus of genotype AT and those which failed to amplify the band given the genotype AA for this locus. The target capture linkage maps constructed for *Triturus* and *Lissotriton* in France et al. (2024a) were then rebuilt to include these calls, thus allowing the location of the Y-linked region on the sequence capture map, otherwise using the same data, settings and pipeline as described in that study. The rebuilt maps were then compared to each other, and the *P. waltl* genome assembly, to highlight any homology between the *Lissotriton* Y-chromosome, and the region associated with the *Triturus* balanced lethal system (or if no sex chromosome turnover has occurred, the *Triturus* Y-chromosome).

## Results

### Sex association

After demultiplexing and initial filtering the known-sex adults yielded a total of 501 million read-pairs (median per sample: 7.62 M, interquartile range: 5.28-9.59 M). A total of 1,394,143 million loci were identified, of which 179,516 (12.88%) were present in at least 50% of all samples. The initial selection for candidate Y-linked markers generated yielded a total of 39 loci.

We designed 23 primer pairs (sup. Table S3) for the 12 highest ranked candidate markers (for one marker TiY-444315, Primer 3 was unable to find a valid primer pair for the shorter fragment). 11 primer pairs, covering 6 marker sequences, amplified products only in male in *T. ivanbureschi*. 6 pairs (for 5 markers) also were male specific in both *T. macedonicus* and *T. cristatus* (Table 1, sup. Fig. S1-3). No primer pairs were successful in all taxa, however TiY-384959-short showed male specificity in all species except *T. dobrogicus*, where no product was amplified in either sex. TiY-137941-long was the only primer pair to amplify in male *T. dobrogicus* and was also male specific in all other taxa that occur within the Balkans (but did not amplify sex-specifically in *T. karelinii*, *T. marmoratus, T. pygmaeus* or the Italian lineage of *T. carnifex*) (Fig. 3, primer sequences in Table 2).

**Figure 3:**
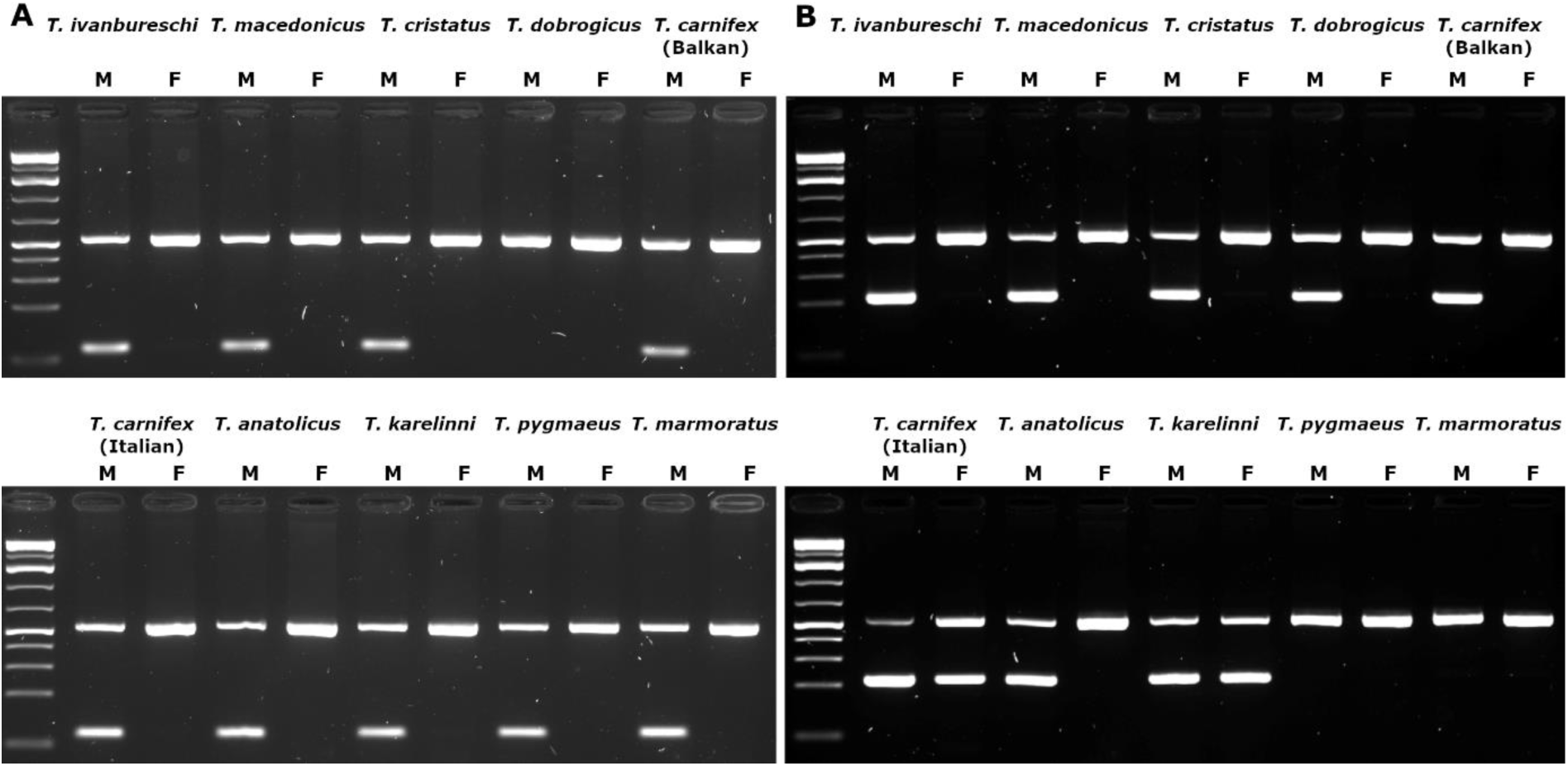
Gels showing amplification of Y-linked primer pairs TiY-384959-short (A) and TiY-137941-long (B), with CDK-17 (537 bp) as control maker, in male-female pairs of various *Triturus* taxa. Male samples are labelled M and females F. TiY-384959-short (92 bp) shows male-specific amplification in all species except *T. dobrogicus*, where no product is seen. TiY-137941-long (ca. 170 bp) is male specific in *T. dobrogicus* and all other taxa which occur in the Balkans (shown on the top row of the gel) but shows no amplification in the marbled newts (*T. marmoratus* and *T. pygmaeus*) and non-sex specific amplification in *T. karelinii* and the Italian lineage of *T. carnifex*.

**Table 1:**
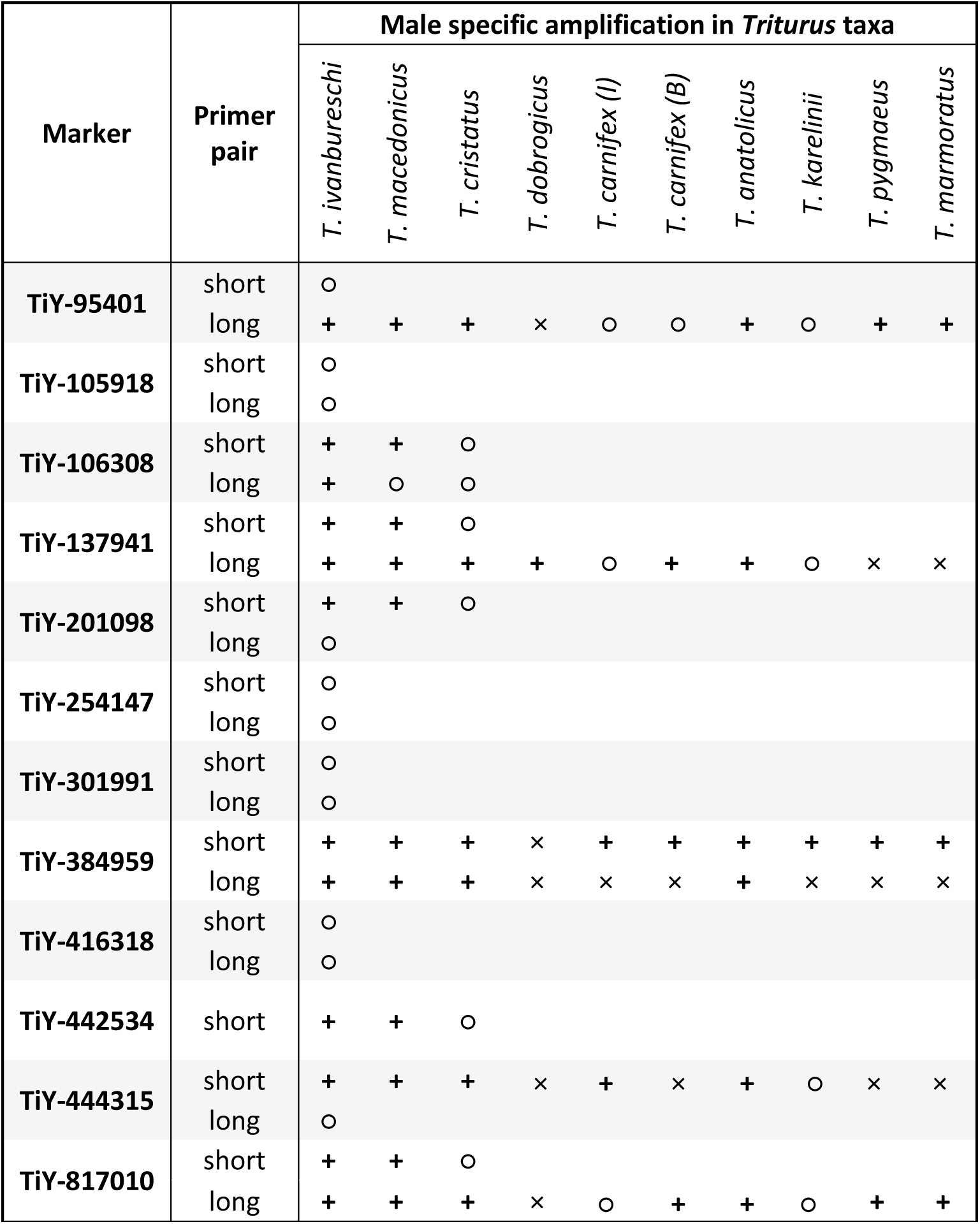
Summary of results of PCR screening of primer pairs designed for candidate Y-linked markers in *Triturus* newts. Results are indicated as: **+** amplification only in male samples, ○ amplification in both male and female samples, × no amplification in either sex. Twenty-three primer pairs were tested in a male-female pair of *T. ivanbureschi*. Twelve primer pairs, covering eight candidate markers, show confirmed male-specific amplification in *T. ivanbureschi*. The successful primer sets were then tested in male-female pairs of *T. macedonicus* and *L. cristatus*, and the six successful in both were then tested in all available *Triturus* taxa. While candidate TiY-384959-short demonstrated broad male-specificity across the genus, it failed to amplify in either male or female *T. dobrogicus* - TiY-137941-long proved the only successful primer pair in this species.

**Table 2:**
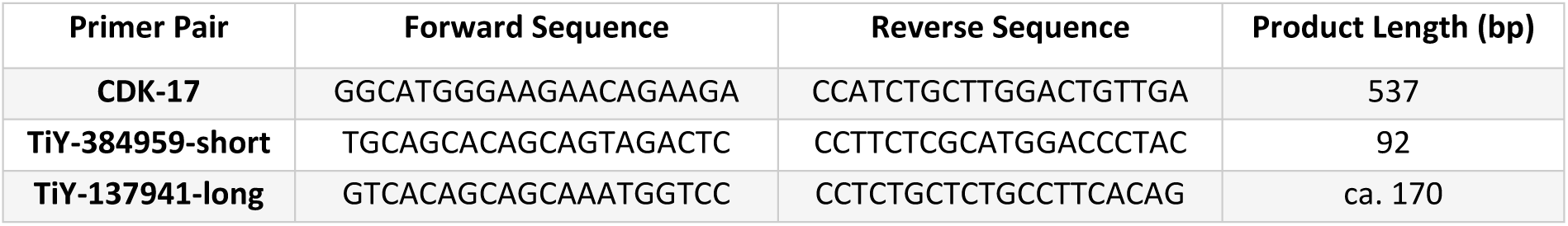
Primer sequences used for the sex diagnostic PCR for use within the genus *Triturus*. CDK-17 is an autosomal control marker. TiY-384959-short is male-specific in all species except *T. dobrogicus*. TiY-137941-long is male-specific in *T. dobrogicus,* as well as *T. ivanbureschi*, *T. anatolicus*, *T. macedonicus*, *T. cristatus* and the Balkan lineage of *T. carnifex*.

### RADseq Linkage map

The linkage map constructed from the *T. ivanbureschi* family consists of 7,233 markers arranged into 12 linkage groups (corresponding to the 12 chromosomes of the *Triturus* genome), with a total length of 1,120 cM (Fig. 4, sup. table S2). Twenty-seven Y-linked presence/absence markers were placed on the map, all located in a 3.2 cM region at the end of linkage group 8, with 24 of these markers being placed at a single point.

**Figure 4:**
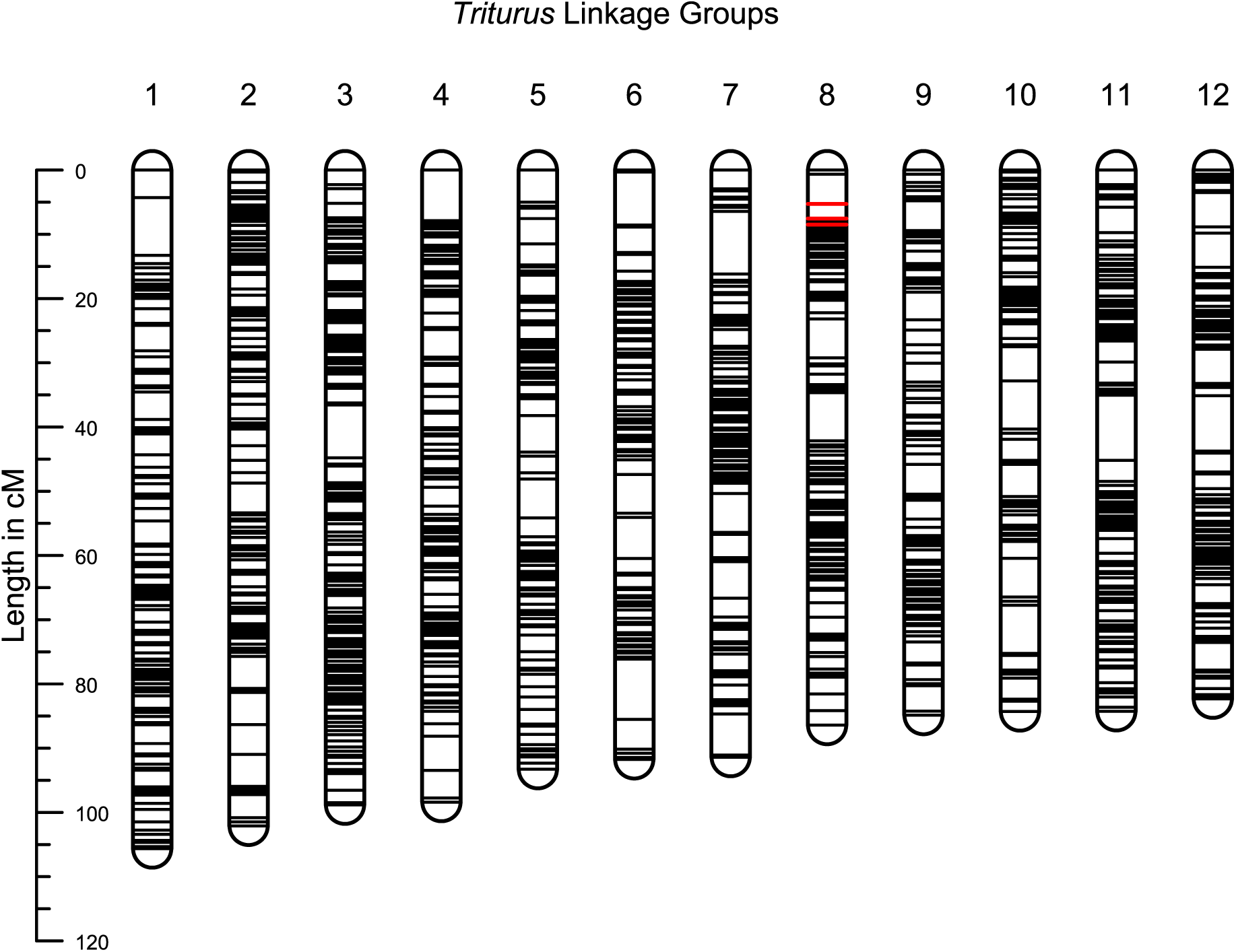
Linkage map for *Triturus ivanbureschi* based on 7,233 RADseq markers, arranged in 12 linkage groups. 27 male-linked presence absence markers – highlighted in red – are located at one end of linkage group 8, identifying it as the Y-chromosome.

Five hundred and twenty-four markers (7.2%) placed on the linkage map, including a single Y-linked marker, could be aligned with sequences from the *P. waltl* genome assembly. Each *T. ivanbureschi* linkage group shows a clear and reciprocal correspondence with one of the *P. waltl* chromosomes, with 374 (71%) markers mapping to their corresponding chromosome, and large-scale synteny within chromosomes. We fail to observe any clear pattern in the remaining markers, indicating that these are likely a consequence of BLAST hits against paralogous sequences, rather than a major rearrangement in the genome of either species.

Forty-seven markers within the *T. ivanbureschi* Y-chromosome (linkage group 8 BLAST against sequences from *P. waltl* chromosome 2 (Fig. 5), including the solitary Y-linked marker. However, in the analogous RADseq linkage map previously made for *L. vulgaris*, the Y-chromosome is clearly seen as homologous to *P. waltl* chromosome 5. We see no evidence that this is a result of translocation of the sex-determining region, no markers from *T. ivanbureschi* linkage group 3 BLAST against sequences from *P. waltl* chromosome 5. A single marker from the *L. vulgaris* Y-chromosome (linkage group 5) is found in *P. waltl* chromosome 2, however this is located over 100 cM away from the sex-linked region.

**Figure 5:**
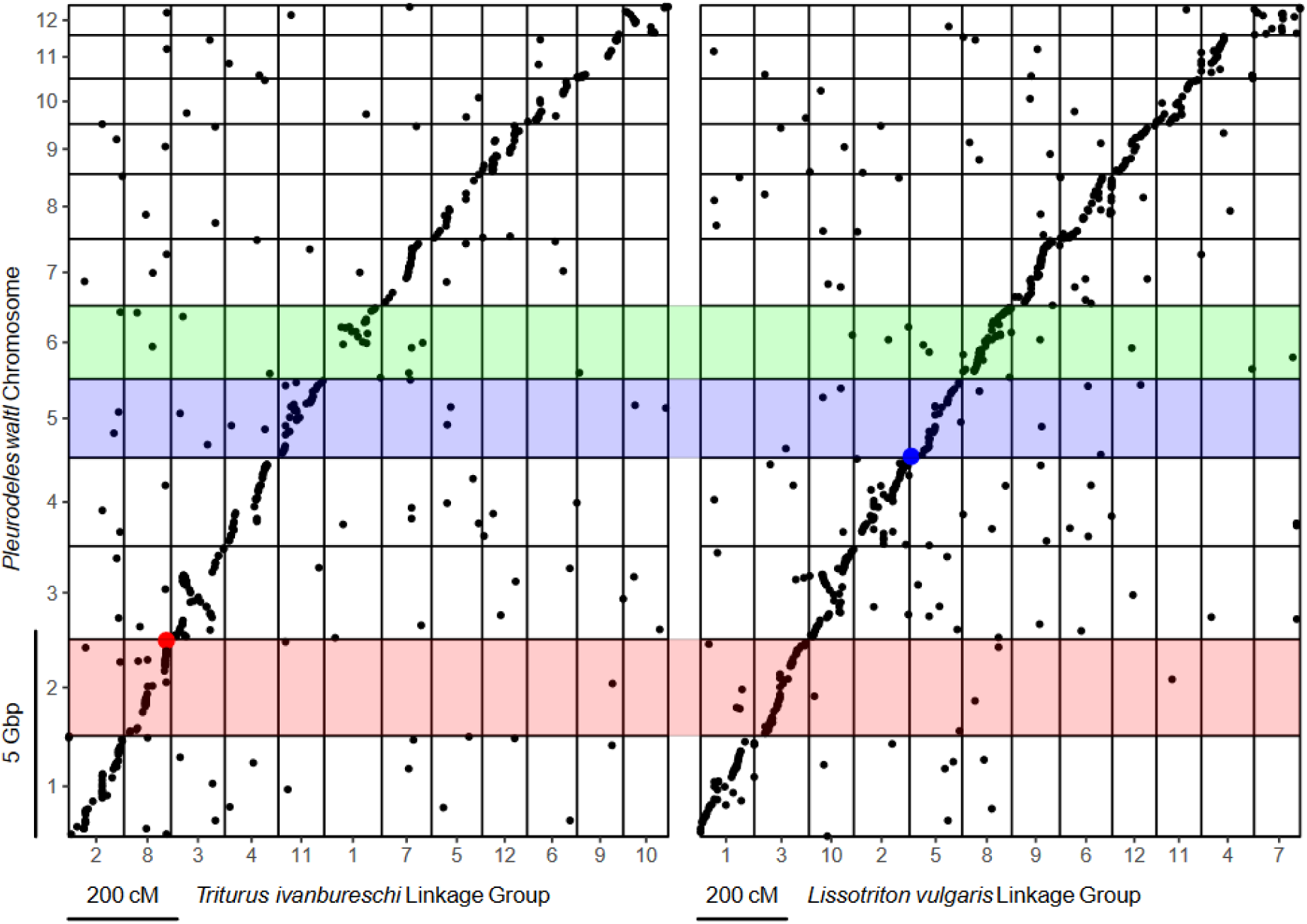
Oxford plots showing the *Triturus ivanbureschi* (left) and *Lissotriton vulgaris* (right – from France et al. (2024b)) RADseq linkage maps against the *Pleurodeles waltl* genome assembly (Brown et al., 2023). The *T. ivanbureschi* Y-linked presence/absence markers identified in known sex-adults are highlighted in red, the *L. vulgaris* Y-linked markers in blue. The *T. ivanbureschi* Y-chromosome is shown to be homologous to *P. waltl* chromosome 2 and thus is not homologous with the *L. vulgaris* Y-chromosome (which is homologous to *P. waltl* chromosome 5). There is no evidence of a translocation of the sex-linked regions of either chromosome. *Pleurodeles waltl* chromosome 6, highlighted in green is homologous to the *Triturus* balanced lethal system on chromosome 1.

### Identification of Y-linked regions in target capture linkage maps

We incorporate Y-linked RAD markers for *Lissotriton* (France et al., 2024b) and *Triturus* (identified in this study) into the target capture-based linkage maps previously constructed for these two genera (France et al., 2024a), which include genes linked to the balanced lethal system. For *Lissotriton*, 110 offspring amplified the Y-linked RAD marker lvY-51393-short, whereas 92 did not, with a single individual giving an ambiguous result (either a missing control band, or only very faint amplification of either band). For *Triturus*, 107 offspring amplified the marker TiY-384959-short, whereas 93 did not – with six giving ambiguous results. Both offspring sets are biased towards males, though this is not statistically significant – assuming an even sex ratio, two-tailed binomial p-values are 0.231 for *Lissotriton* and 0.358 for *Triturus*.

For both *Triturus* and *Lissotriton* the number of genotype calls is sufficient to confidently locate the Y-linked RAD markers within the target capture linkage maps (Fig. 6). In concordance with the RADseq linkage maps the *Triturus* and *Lissotriton* Y-chromosomes are shown not to be homologous, with the *Triturus* and *Lissotriton* Y-linked regions again located on the homologs of *P. waltl* chromosomes 2 and 5. However, the *Lissotriton* Y-chromosome also lacks homology with *Triturus* chromosome 1, where the genes associated with the balanced lethal system are located (Fig. 6). Instead *Triturus* chromosome 1 is homologous with *P. waltl* chromosome 6.

**Figure 6:**
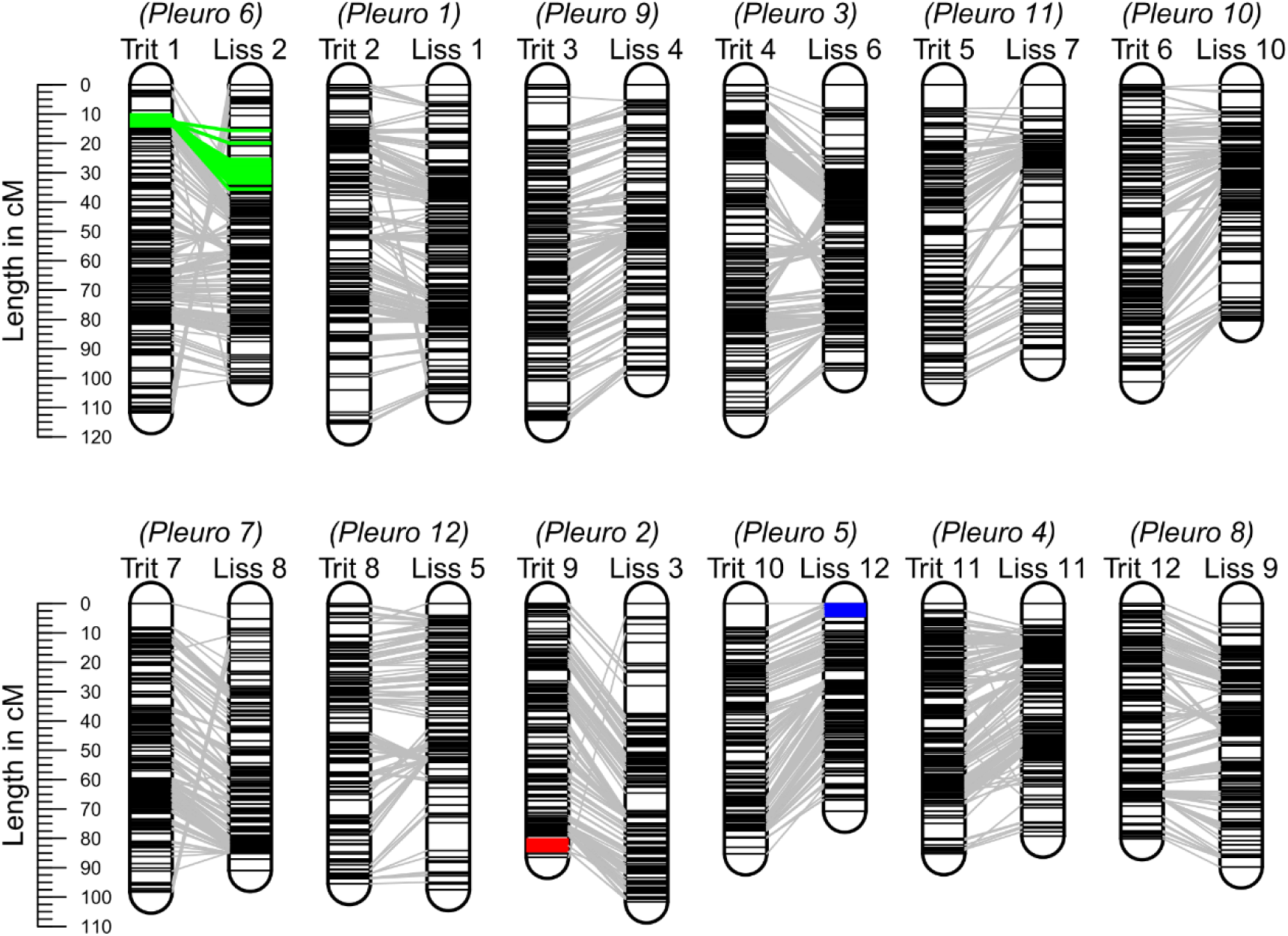
Target capture linkage maps (France et al., 2024a) augmented with Y-linked markers (the abbreviations Trit and Liss refer to the linkage groups from *Triturus* and *Lissotriton* respectively – Pleuro refers to the homologous chromosome in the *Pleurodeles waltl* genome assembly). *Triturus* balanced lethal system associated markers are highlighted in green, the *Triturus* Y-linked marker in red and the *Lissotriton* Y-linked marker in blue. In accordance with the RADseq linkage maps, the *Triturus* and *Lissotriton* Y-chromosomes are shown not to be homologous. Additionally, the *Lissotriton* Y-chromosome is also not homologous with *Triturus* chromosome 1. Note that the numbering of the linkage groups differs between the RADseq and target capture linkage maps.

## Discussion

### No homology between the balanced lethal system and sex chromosomes

The hypothesis that the *Triturus* balanced lethal system evolved from a sex chromosome system (Grossen et al., 2012) implies a pair of predictions that we test in this study. Firstly, that sex chromosome turnover must have occurred in *Triturus* after it diverged from its sister lineage *Lissotriton* and so these taxa cannot now share a sex-determination system. Secondly, that *Triturus* chromosome 1 must be homologous to the sex chromosome of this common ancestor, and so would be homologous to the modern *Lissotriton* Y-chromosome – unless this genus has also undergone sex chromosome turnover.

We confidently identify the *Triturus* Y-chromosome by identifying a set of male-linked markers and placing them within both a newly constructed high density RADseq linkage map, and a previously constructed map based on target capture. We discover that the *Triturus* Y-chromosome is clearly not homologous to the *Lissotriton* Y-chromosome. Therefore, at least one of these lineages must have undergone a sex chromosome turnover, which is compatible with the first prediction. However, this could also be explained by sex chromosome turnover within the *Lissotriton* lineage, and as no sex-linked markers are known for any other newt taxa, we lack an outgroup to distinguish between these scenarios.

We show that, counter to the second prediction made by the sex chromosome origin hypothesis, the *L. vulgaris* Y-linked region is clearly not homologous to the balanced lethal system present on *Triturus* chromosome 1. Although we cannot rule out the possibility that sex chromosome turnover has occurred in both *Lissotriton* and *Triturus*, such that neither taxon now possesses the ancestral Y-chromosome, the most parsimonious explanation of our results is a single sex chromosome turnover after the divergence of the *Triturus* and *Lissotriton* lineages. If the *Triturus* balanced lethal system did arise from the ancestral sex chromosome, this would require at least one additional turnover. The plausibility of the sex chromosome origin hypothesis thus depends on how common sex chromosome turnover events are within newts.

### Y-chromosome switching in salamanders: common or rare?

Unlike mammals and birds (Cortez et al., 2014; Ellegren, 2010), sex chromosome turnover appears relatively common in amphibians (Miura, 2017). However, within salamanders, the large genome size and consequent difficulty in discovering sex-linked sequences has meant that its observation has only been possible in cases where female heterogametery (ZW) has transitioned to male heterogametery (XY) or vice versa – at least three such events are known (Hime et al., 2019). Conversely, little is known about the frequency of transitions between different Y-linked (or W-linked) sex determination systems in the salamanders. There is some evidence from other amphibians that these events may be common. In the family Ranidae (true frogs), Jeffries et al. (2018) found 13 sex chromosome turnover events in 28 lineages, 11 of which were transitions between different Y-chromosomes. Additionally, the X and Y-chromosomes in *Triturus, Lissotriton,* and other related newt genera such as *Ichthyosaura* are poorly differentiated (Schmid et al., 1979; Sims et al., 1984), which may be taken as evidence that they have all evolved rather recently (Charlesworth & Charlesworth, 2000).

Nonetheless, we also have evidence of sex chromosome stasis in salamanders. Despite having diverged in late Cretaceous, the giant salamanders (the family Cryptobranchidae), possess a conserved W-linked region, with the same female specific marker shown to amplify in both the North American hellbender and the Chinese giant salamander (Hime et al., 2019). These Z and W-chromosomes also appear extremely homomorphic (Sessions et al., 1982), showing that this is not necessarily proof of a recent origin. A similar phenomenon of deceptively youthful sex chromosomes is also seen in tree frogs of the genus *Hyla* (Stöck et al., 2011). At present there is insufficient data to determine whether the Y-to-Y-chromosome turnover we observe between *Triturus* and *Lissotriton* is a common or exceptional event. The identification of sex determining regions in other salamander genera would help to answer this question.

### Towards sex chromosome identification across salamanders

The sex associative RADseq methodology we employ in this study is a relatively quick and effective approach for the discovery of sex-linked sequences. These markers are not just useful for evolutionary genomics, but also invaluable for researchers interested in the ecology or population dynamics, especially in species which are difficult to morphologically sex before maturity, such as newts (Sparreboom, 2014). In the case of *Triturus* we can recommend the marker TiY-384959-short in any context except when the Danube crested newt, *T. dobrogicus* may be encountered, where TiY-137941-long should be used instead.

However, while the identification of sex-linked markers is simple, determining whether they are homologous often requires locating them within a genome. Whether by linkage mapping, whole genome assembly or methods such as fluorescent in-situ hybridisation, this is often a resource intensive process. The need for such investment may be circumnavigated by aligning Y-linked RAD sequences with a genome from a related organism. In our study we show some success by employing the genome of the relatively distantly related *Pleurodeles waltl* which diverged over 60 mya (Marjanović & Laurin, 2014), although only one of 27 Y-linked markers could be confidently aligned. For the investigation of newt Y-chromosomes specifically, whole genome data from a more closely related species would be extremely valuable. Given the high degree of chromosome level synteny we observe between *P. waltl*, *Triturus*, and *Lissotriton* we suggest that simply scaffolding long read data against the *P. waltl* assembly would result in a reference genome sufficient for locating sex-linked regions, similar to the strategy employed by Jeffries et al. (2018) in *Rana*. Additionally, such long read data could be used to identify sequences more conserved than those derived directly from RADseq, allowing for the development of markers that show sex-specific amplification in multiple genera.

Further investigation of Y-linked markers in newts thus promises insights into the rate of sex chromosome turnover in salamanders, as well as determining which, if either, of the *Triturus* and *Lissotriton* Y-chromosomes are ancestral – and if a sex chromosome turnover is an at all plausible explanation of the *Triturus* balanced lethal system.

## Acknowledgements

This project has received funding from the European Research Council (ERC) under the European Union’s Horizon 2020 research and innovation programme (Grant Agreement No. 802759). Tissue samples for adult *T. ivanbureschi* of known sex were taken from Zli Dol, Serbia (permit no. 353-01-75/2014-08 of the Ministry of Energy, Development and Environmental Protection). The experimental procedures to breed the linkage map family were approved by the Ethics Committee of the Institute for Biological Research „Siniša Stanković”, University of Belgrade (decisions no. 03-03/16 and 01-1949). Experiments were performed in accordance with the Directive 2010/63/EU and supported by the Serbian Ministry of Science, Technological Development and Innovation (grants no. 451-03-65/2024-03/ 200178 and 451-03-66/2024-03/ 200178). We give thanks to Dr. Christine Grossen for her feedback on a draft of this manuscript.

## Data Availability

All raw reads can be found as a part of the NCBI accession associated with Bioproject: PRJNA1173742. (https://www.ncbi.nlm.nih.gov/bioproject/PRJNA1173742). The scripts and bioinformatic pipelines used for analysis are available at the GitHub repository: Wielstra-Lab/Triturus_RADseq_Y (https://github.com/Wielstra-Lab/Triturus_RADseq_Y). The positions and sequences of all markers in the Triturus RADseq linkage map can be found in a .xlsx file hosted in the Zenodo repository with DOI 10.5281/zenodo.14288865.

