## Supplementary Data for "Identification of Y-chromosome turnover in newts fails to support a sex chromosome origin for the *Triturus* balanced lethal system"

**Table of Contents:**


| | |
|---|---|
| **Figure S1: Screening in *T. ivanbureschi*** | Page 2 |
| **Figure S2: Screening in *T. cristatus* and *T. macedonicus*** | Page 2 |
| **Figure S3: Screening in other *Triturus* species** | Page 3 |
| **Table S1: Sample information** | Page 4 |
| **Table S2: Linkage map statistics** | Page 7 |
| **Table S3: Primer sequences** | Page 7 |

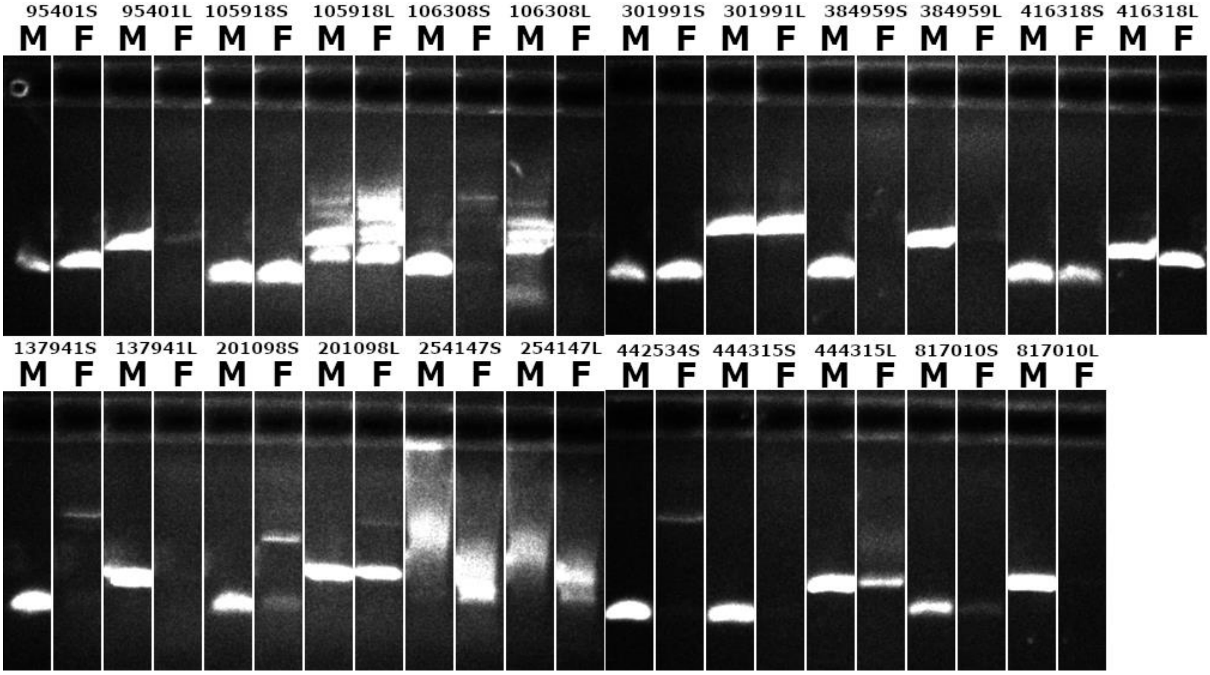


**Figure S1:** PCR screening of 23 primer pairs designed for candidate Y-linked markers for sex specific amplification in *Triturus ivanbureschi*. Label M indicates the male sample and label F indicates female. Markers are indicated by number followed by either S (for primer pairs designed for the short product – c.a. 100 bp) or L (for primer pairs designed for the long product – c.a. 200 bp). 12 pairs show male specific amplification.

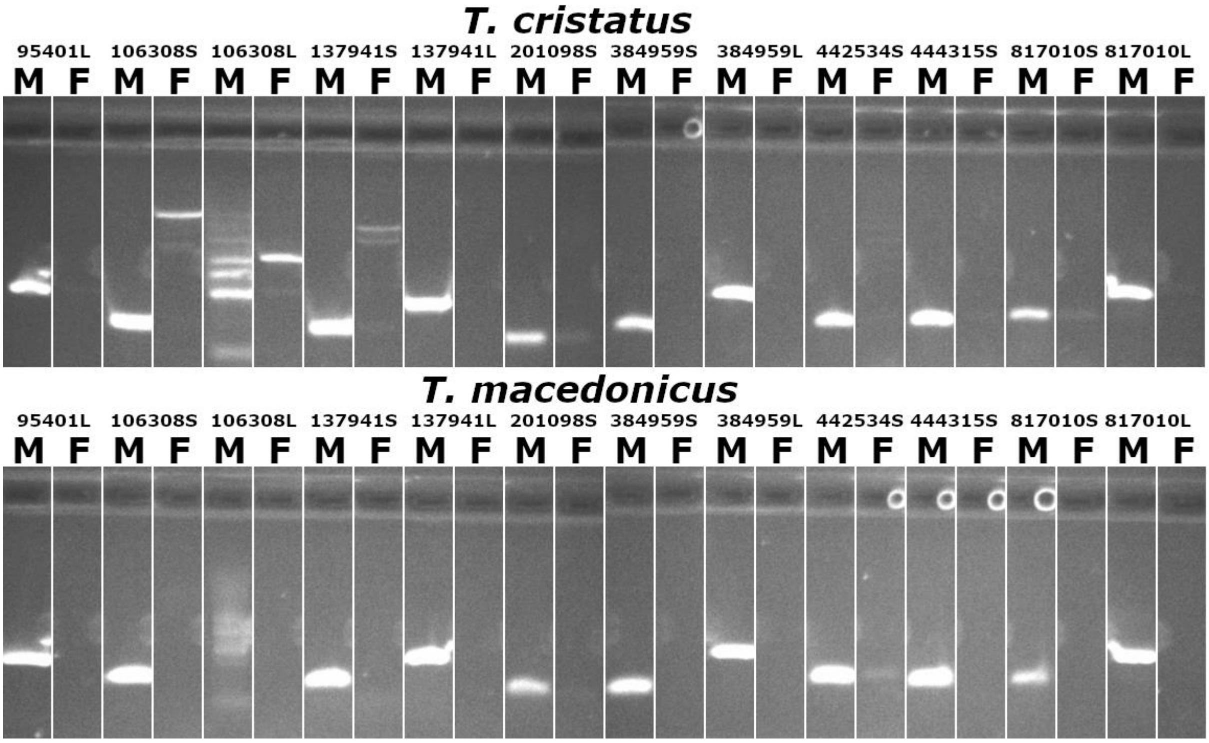


**Figure S2:** Further PCR screening of 12 primer pairs for Y-linked markers (that show male specific amplification in *T. ivanbureschi*) in *T. cristatus* and *T. macedonicus*. Label M indicates the male sample and label F indicates female. Markers are indicated by number followed by either S (for primer pairs designed for the short product – c.a. 100 bp) or L (for primer pairs designed for the long product – c.a. 200 bp). 6 primer pairs show strong amplification in males of both species with no product at all visible in females (several other show varying degrees of weak amplification in females).

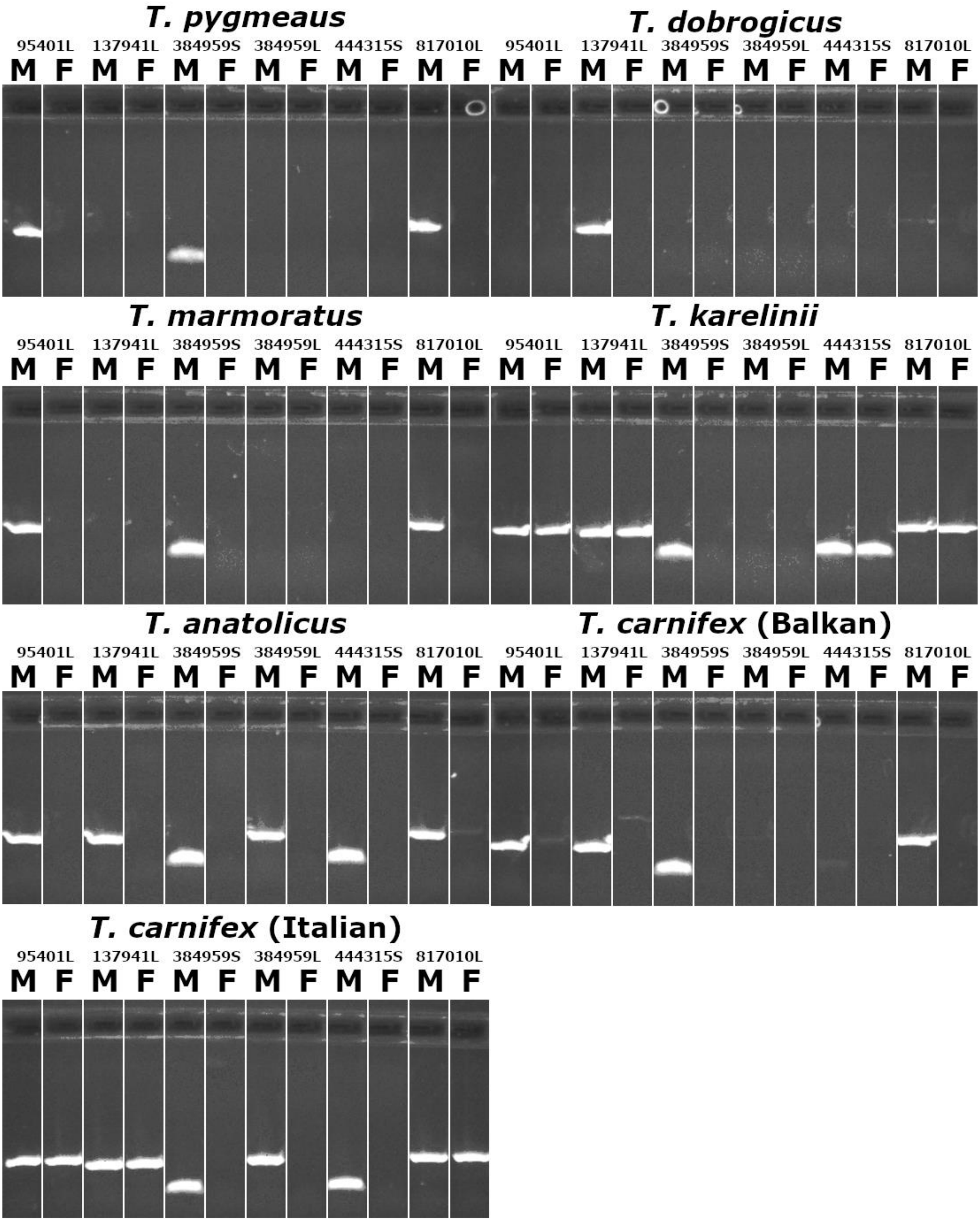


**Figure S3:** PCR screening of 6 primer pairs for Y-linked markers (that show confirmed male specific amplification in three *Triturus* species) in all other *Triturus* species (except for *T. rudolfi*, which was not yet described at the time of study). Label M indicates the male sample and label F indicates female. Markers are indicated by number followed by either S (for primer pairs designed for the short product – c.a. 100 bp) or L (for primer pairs designed for the long product – c.a. 200 bp). No marker shows male specific amplification in all species, however TiY-384959-short is successful in all other than *T. dobrogicus*. The only marker to show any amplification in *T. dobrogicus* is TiY-137941-long.

**Table S1:** Samples used within this study. All samples designated for use as 'Sexed RADseq' and 'Map parent' were collected in Zli Dol (Pčinja district, Serbia, 42°25 N; 22°27 E). All other samples were captive bred. RADseq data from all *Triturus ivanbureschi* samples is available as part of NCBI bioproject PRJNA1173742.

| Sample | Species | Sex | Use | Sample | Species | Sex | Use |
|---|---|---|---|---|---|---|---|
| **BW_0008** | *T. ivanbureschi* | Male | Map parent | **BW_0576** | *T. ivanbureschi* | Unknown | Map offspring |
| **BW_0009** | *T. ivanbureschi* | Female | Map parent | **BW_0577** | *T. ivanbureschi* | Unknown | Map offspring |
| **BW_0798** | *T. cristatus* | Male | Screening | **BW_0578** | *T. ivanbureschi* | Unknown | Map offspring |
| **BW_0799** | *T. cristatus* | Female | Screening | **BW_0579** | *T. ivanbureschi* | Unknown | Map offspring |
| **BW_0803** | *T. macedonicus* | Male | Screening | **BW_0580** | *T. ivanbureschi* | Unknown | Map offspring |
| **BW_0804** | *T. macedonicus* | Female | Screening | **BW_0581** | *T. ivanbureschi* | Unknown | Map offspring |
| **BW_0786** | *T. Pygmaeus* | Male | Screening | **BW_0585** | *T. ivanbureschi* | Unknown | Map offspring |
| **BW_0787** | *T. Pygmaeus* | Female | Screening | **BW_0587** | *T. ivanbureschi* | Unknown | Map offspring |
| **BW_0788** | *T. Dobrogicus* | Male | Screening | **BW_0588** | *T. ivanbureschi* | Unknown | Map offspring |
| **BW_0789** | *T. Dobrogicus* | Female | Screening | **BW_0590** | *T. ivanbureschi* | Unknown | Map offspring |
| **BW_0790** | *T. Marmoratus* | Male | Screening | **BW_0592** | *T. ivanbureschi* | Unknown | Map offspring |
| **BW_0791** | *T. Marmoratus* | Female | Screening | **BW_0594** | *T. ivanbureschi* | Unknown | Map offspring |
| **BW_0792** | *T. Karelinii* | Male | Screening | **BW_0595** | *T. ivanbureschi* | Unknown | Map offspring |
| **BW_0793** | *T. Karelinii* | Female | Screening | **BW_0597** | *T. ivanbureschi* | Unknown | Map offspring |
| **BW_0796** | *T. Anatolicus* | Male | Screening | **BW_0599** | *T. ivanbureschi* | Unknown | Map offspring |
| **BW_0797** | *T. Anatolicus* | Female | Screening | **BW_0601** | *T. ivanbureschi* | Unknown | Map offspring |
| **BW_0800** | *T. Carnifex (B)* | Male | Screening | **BW_0602** | *T. ivanbureschi* | Unknown | Map offspring |
| **BW_0801** | *T. Carnifex (B)* | Female | Screening | **BW_0605** | *T. ivanbureschi* | Unknown | Map offspring |
| **BW_0805** | *T. Carnifex (I)* | Male | Screening | **BW_0606** | *T. ivanbureschi* | Unknown | Map offspring |
| **BW_0806** | *T. Carnifex (I)* | Female | Screening | **BW_0607** | *T. ivanbureschi* | Unknown | Map offspring |
| **BW_0758** | *T. ivanbureschi* | Male | Sexed RADseq | **BW_0609** | *T. ivanbureschi* | Unknown | Map offspring |
| **BW_0759** | *T. ivanbureschi* | Male | Sexed RADseq | **BW_0611** | *T. ivanbureschi* | Unknown | Map offspring |
| **BW_0761** | *T. ivanbureschi* | Male | Sexed RADseq | **BW_0612** | *T. ivanbureschi* | Unknown | Map offspring |
| **BW_0762** | *T. ivanbureschi* | Male | Sexed RADseq | **BW_0615** | *T. ivanbureschi* | Unknown | Map offspring |
| **BW_0763** | *T. ivanbureschi* | Male | Sexed RADseq | **BW_0617** | *T. ivanbureschi* | Unknown | Map offspring |
| **BW_0764** | *T. ivanbureschi* | Male | Sexed RADseq | **BW_0618** | *T. ivanbureschi* | Unknown | Map offspring |
| **BW_0765** | *T. ivanbureschi* | Male | Sexed RADseq | **BW_0619** | *T. ivanbureschi* | Unknown | Map offspring |
| **BW_0766** | *T. ivanbureschi* | Male | Sexed RADseq | **BW_0620** | *T. ivanbureschi* | Unknown | Map offspring |
| **BW_0767** | *T. ivanbureschi* | Male | Sexed RADseq | **BW_0621** | *T. ivanbureschi* | Unknown | Map offspring |
| **BW_0768** | *T. ivanbureschi* | Male | Sexed RADseq | **BW_0623** | *T. ivanbureschi* | Unknown | Map offspring |
| **BW_0769** | *T. ivanbureschi* | Male | Sexed RADseq | **BW_0625** | *T. ivanbureschi* | Unknown | Map offspring |
| **BW_0770** | *T. ivanbureschi* | Male | Sexed RADseq | **BW_0626** | *T. ivanbureschi* | Unknown | Map offspring |
| **BW_0771** | *T. ivanbureschi* | Male | Sexed RADseq | **BW_0628** | *T. ivanbureschi* | Unknown | Map offspring |
| **BW_0772** | *T. ivanbureschi* | Female | Sexed RADseq | **BW_0631** | *T. ivanbureschi* | Unknown | Map offspring |
| **BW_0773** | *T. ivanbureschi* | Female | Sexed RADseq | **BW_0633** | *T. ivanbureschi* | Unknown | Map offspring |
| **BW_0774** | *T. ivanbureschi* | Female | Sexed RADseq | **BW_0634** | *T. ivanbureschi* | Unknown | Map offspring |
| **BW_0775** | *T. ivanbureschi* | Female | Sexed RADseq | **BW_0635** | *T. ivanbureschi* | Unknown | Map offspring |
| **BW_0776** | *T. ivanbureschi* | Female | Sexed RADseq | **BW_0636** | *T. ivanbureschi* | Unknown | Map offspring |
| **BW_0777** | *T. ivanbureschi* | Female | Sexed RADseq | **BW_0639** | *T. ivanbureschi* | Unknown | Map offspring |
| **BW_0778** | *T. ivanbureschi* | Female | Sexed RADseq | **BW_0644** | *T. ivanbureschi* | Unknown | Map offspring |
| **BW_0780** | *T. ivanbureschi* | Female | Sexed RADseq | **BW_0645** | *T. ivanbureschi* | Unknown | Map offspring |
| **BW_0781** | *T. ivanbureschi* | Female | Sexed RADseq | **BW_0646** | *T. ivanbureschi* | Unknown | Map offspring |
| **BW_0782** | *T. ivanbureschi* | Female | Sexed RADseq | **BW_0648** | *T. ivanbureschi* | Unknown | Map offspring |

| Sample | Species | Sex | Use |
|---|---|---|---|
| **BW_0783** | *T. ivanbureschi* | Female | Sexed RADseq |
| **BW_0784** | *T. ivanbureschi* | Female | Sexed RADseq |
| **BW_0785** | *T. ivanbureschi* | Female | Sexed RADseq |
| **BW_2095** | *T. ivanbureschi* | Male | Sexed RADseq |
| **BW_2096** | *T. ivanbureschi* | Male | Sexed RADseq |
| **BW_2097** | *T. ivanbureschi* | Male | Sexed RADseq |
| **BW_2098** | *T. ivanbureschi* | Male | Sexed RADseq |
| **BW_2099** | *T. ivanbureschi* | Male | Sexed RADseq |
| **BW_2100** | *T. ivanbureschi* | Male | Sexed RADseq |
| **BW_2101** | *T. ivanbureschi* | Male | Sexed RADseq |
| **BW_2102** | *T. ivanbureschi* | Male | Sexed RADseq |
| **BW_2103** | *T. ivanbureschi* | Male | Sexed RADseq |
| **BW_2104** | *T. ivanbureschi* | Male | Sexed RADseq |
| **BW_2105** | *T. ivanbureschi* | Male | Sexed RADseq |
| **BW_2106** | *T. ivanbureschi* | Male | Sexed RADseq |
| **BW_2107** | *T. ivanbureschi* | Male | Sexed RADseq |
| **BW_2108** | *T. ivanbureschi* | Male | Sexed RADseq |
| **BW_2110** | *T. ivanbureschi* | Male | Sexed RADseq |
| **BW_2111** | *T. ivanbureschi* | Female | Sexed RADseq |
| **BW_2112** | *T. ivanbureschi* | Female | Sexed RADseq |
| **BW_2113** | *T. ivanbureschi* | Female | Sexed RADseq |
| **BW_2114** | *T. ivanbureschi* | Female | Sexed RADseq |
| **BW_2115** | *T. ivanbureschi* | Female | Sexed RADseq |
| **BW_2116** | *T. ivanbureschi* | Female | Sexed RADseq |
| **BW_2117** | *T. ivanbureschi* | Female | Sexed RADseq |
| **BW_2118** | *T. ivanbureschi* | Female | Sexed RADseq |
| **BW_2119** | *T. ivanbureschi* | Female | Sexed RADseq |
| **BW_2120** | *T. ivanbureschi* | Female | Sexed RADseq |
| **BW_2121** | *T. ivanbureschi* | Female | Sexed RADseq |
| **BW_2122** | *T. ivanbureschi* | Female | Sexed RADseq |
| **BW_2123** | *T. ivanbureschi* | Female | Sexed RADseq |
| **BW_2124** | *T. ivanbureschi* | Female | Sexed RADseq |
| **BW_2125** | *T. ivanbureschi* | Female | Sexed RADseq |
| **BW_2126** | *T. ivanbureschi* | Female | Sexed RADseq |
| **BW_2127** | *T. ivanbureschi* | Male | Sexed RADseq |
| **BW_0536** | *T. ivanbureschi* | Unknown | Map offspring |
| **BW_0537** | *T. ivanbureschi* | Unknown | Map offspring |
| **BW_0538** | *T. ivanbureschi* | Unknown | Map offspring |
| **BW_0539** | *T. ivanbureschi* | Unknown | Map offspring |
| **BW_0540** | *T. ivanbureschi* | Unknown | Map offspring |
| **BW_0541** | *T. ivanbureschi* | Unknown | Map offspring |
| **BW_0542** | *T. ivanbureschi* | Unknown | Map offspring |
| **BW_0543** | *T. ivanbureschi* | Unknown | Map offspring |
| **BW_0544** | *T. ivanbureschi* | Unknown | Map offspring |
| **BW_0545** | *T. ivanbureschi* | Unknown | Map offspring |
| **BW_0546** | *T. ivanbureschi* | Unknown | Map offspring |
| **BW_0650** | *T. ivanbureschi* | Unknown | Map offspring |
| **BW_0653** | *T. ivanbureschi* | Unknown | Map offspring |
| **BW_0655** | *T. ivanbureschi* | Unknown | Map offspring |
| **BW_0657** | *T. ivanbureschi* | Unknown | Map offspring |
| **BW_0659** | *T. ivanbureschi* | Unknown | Map offspring |
| **BW_0660** | *T. ivanbureschi* | Unknown | Map offspring |
| **BW_0661** | *T. ivanbureschi* | Unknown | Map offspring |
| **BW_0664** | *T. ivanbureschi* | Unknown | Map offspring |
| **BW_0665** | *T. ivanbureschi* | Unknown | Map offspring |
| **BW_0666** | *T. ivanbureschi* | Unknown | Map offspring |
| **BW_0667** | *T. ivanbureschi* | Unknown | Map offspring |
| **BW_0668** | *T. ivanbureschi* | Unknown | Map offspring |
| **BW_0670** | *T. ivanbureschi* | Unknown | Map offspring |
| **BW_0671** | *T. ivanbureschi* | Unknown | Map offspring |
| **BW_0673** | *T. ivanbureschi* | Unknown | Map offspring |
| **BW_0674** | *T. ivanbureschi* | Unknown | Map offspring |
| **BW_0676** | *T. ivanbureschi* | Unknown | Map offspring |
| **BW_0677** | *T. ivanbureschi* | Unknown | Map offspring |
| **BW_0678** | *T. ivanbureschi* | Unknown | Map offspring |
| **BW_0679** | *T. ivanbureschi* | Unknown | Map offspring |
| **BW_0680** | *T. ivanbureschi* | Unknown | Map offspring |
| **BW_0682** | *T. ivanbureschi* | Unknown | Map offspring |
| **BW_0683** | *T. ivanbureschi* | Unknown | Map offspring |
| **BW_0684** | *T. ivanbureschi* | Unknown | Map offspring |
| **BW_0685** | *T. ivanbureschi* | Unknown | Map offspring |
| **BW_0686** | *T. ivanbureschi* | Unknown | Map offspring |
| **BW_0689** | *T. ivanbureschi* | Unknown | Map offspring |
| **BW_0690** | *T. ivanbureschi* | Unknown | Map offspring |
| **BW_0691** | *T. ivanbureschi* | Unknown | Map offspring |
| **BW_0692** | *T. ivanbureschi* | Unknown | Map offspring |
| **BW_0693** | *T. ivanbureschi* | Unknown | Map offspring |
| **BW_0694** | *T. ivanbureschi* | Unknown | Map offspring |
| **BW_0695** | *T. ivanbureschi* | Unknown | Map offspring |
| **BW_0697** | *T. ivanbureschi* | Unknown | Map offspring |
| **BW_0698** | *T. ivanbureschi* | Unknown | Map offspring |
| **BW_0699** | *T. ivanbureschi* | Unknown | Map offspring |
| **BW_0701** | *T. ivanbureschi* | Unknown | Map offspring |
| **BW_0702** | *T. ivanbureschi* | Unknown | Map offspring |
| **BW_0703** | *T. ivanbureschi* | Unknown | Map offspring |
| **BW_0704** | *T. ivanbureschi* | Unknown | Map offspring |
| **BW_0705** | *T. ivanbureschi* | Unknown | Map offspring |
| **BW_0706** | *T. ivanbureschi* | Unknown | Map offspring |
| **BW_0707** | *T. ivanbureschi* | Unknown | Map offspring |
| **BW_0708** | *T. ivanbureschi* | Unknown | Map offspring |
| **BW_0709** | *T. ivanbureschi* | Unknown | Map offspring |
| **BW_0710** | *T. ivanbureschi* | Unknown | Map offspring |

| Sample | Species | Sex | Use |
|---|---|---|---|
| **BW_0547** | *T. ivanbureschi* | Unknown | Map offspring |
| **BW_0548** | *T. ivanbureschi* | Unknown | Map offspring |
| **BW_0549** | *T. ivanbureschi* | Unknown | Map offspring |
| **BW_0550** | *T. ivanbureschi* | Unknown | Map offspring |
| **BW_0551** | *T. ivanbureschi* | Unknown | Map offspring |
| **BW_0552** | *T. ivanbureschi* | Unknown | Map offspring |
| **BW_0553** | *T. ivanbureschi* | Unknown | Map offspring |
| **BW_0554** | *T. ivanbureschi* | Unknown | Map offspring |
| **BW_0555** | *T. ivanbureschi* | Unknown | Map offspring |
| **BW_0556** | *T. ivanbureschi* | Unknown | Map offspring |
| **BW_0557** | *T. ivanbureschi* | Unknown | Map offspring |
| **BW_0558** | *T. ivanbureschi* | Unknown | Map offspring |
| **BW_0559** | *T. ivanbureschi* | Unknown | Map offspring |
| **BW_0560** | *T. ivanbureschi* | Unknown | Map offspring |
| **BW_0561** | *T. ivanbureschi* | Unknown | Map offspring |
| **BW_0562** | *T. ivanbureschi* | Unknown | Map offspring |
| **BW_0563** | *T. ivanbureschi* | Unknown | Map offspring |
| **BW_0564** | *T. ivanbureschi* | Unknown | Map offspring |
| **BW_0565** | *T. ivanbureschi* | Unknown | Map offspring |
| **BW_0566** | *T. ivanbureschi* | Unknown | Map offspring |
| **BW_0567** | *T. ivanbureschi* | Unknown | Map offspring |
| **BW_0568** | *T. ivanbureschi* | Unknown | Map offspring |
| **BW_0569** | *T. ivanbureschi* | Unknown | Map offspring |
| **BW_0570** | *T. ivanbureschi* | Unknown | Map offspring |
| **BW_0571** | *T. ivanbureschi* | Unknown | Map offspring |
| **BW_0572** | *T. ivanbureschi* | Unknown | Map offspring |
| **BW_0573** | *T. ivanbureschi* | Unknown | Map offspring |
| **BW_0574** | *T. ivanbureschi* | Unknown | Map offspring |
| **BW_0575** | *T. ivanbureschi* | Unknown | Map offspring |

| Sample | Species | Sex | Use |
|---|---|---|---|
| **BW_0711** | *T. ivanbureschi* | Unknown | Map offspring |
| **BW_0712** | *T. ivanbureschi* | Unknown | Map offspring |
| **BW_0713** | *T. ivanbureschi* | Unknown | Map offspring |
| **BW_0715** | *T. ivanbureschi* | Unknown | Map offspring |
| **BW_0717** | *T. ivanbureschi* | Unknown | Map offspring |
| **BW_0719** | *T. ivanbureschi* | Unknown | Map offspring |
| **BW_0720** | *T. ivanbureschi* | Unknown | Map offspring |
| **BW_0722** | *T. ivanbureschi* | Unknown | Map offspring |
| **BW_0724** | *T. ivanbureschi* | Unknown | Map offspring |
| **BW_0725** | *T. ivanbureschi* | Unknown | Map offspring |
| **BW_0726** | *T. ivanbureschi* | Unknown | Map offspring |
| **BW_0727** | *T. ivanbureschi* | Unknown | Map offspring |
| **BW_0728** | *T. ivanbureschi* | Unknown | Map offspring |
| **BW_0733** | *T. ivanbureschi* | Unknown | Map offspring |
| **BW_0735** | *T. ivanbureschi* | Unknown | Map offspring |
| **BW_0737** | *T. ivanbureschi* | Unknown | Map offspring |
| **BW_0740** | *T. ivanbureschi* | Unknown | Map offspring |
| **BW_0741** | *T. ivanbureschi* | Unknown | Map offspring |
| **BW_0744** | *T. ivanbureschi* | Unknown | Map offspring |
| **BW_0745** | *T. ivanbureschi* | Unknown | Map offspring |
| **BW_0746** | *T. ivanbureschi* | Unknown | Map offspring |
| **BW_0748** | *T. ivanbureschi* | Unknown | Map offspring |
| **BW_0749** | *T. ivanbureschi* | Unknown | Map offspring |
| **BW_0750** | *T. ivanbureschi* | Unknown | Map offspring |
| **BW_0751** | *T. ivanbureschi* | Unknown | Map offspring |
| **BW_0752** | *T. ivanbureschi* | Unknown | Map offspring |
| **BW_0753** | *T. ivanbureschi* | Unknown | Map offspring |
| **BW_0754** | *T. ivanbureschi* | Unknown | Map offspring |
| **BW_0755** | *T. ivanbureschi* | Unknown | Map offspring |

| Group | Number of Markers | Length (cM) |
|---|---|---|
| 1 | 391 | 105.6 |
| 2 | 931 | 102.1 |
| 3 | 1005 | 98.8 |
| 4 | 764 | 98.4 |
| 5 | 331 | 93.8 |
| 6 | 427 | 93.3 |
| 7 | 381 | 91.7 |
| 8 | 600 | 91.4 |
| 9 | 710 | 91.4 |
| 10 | 903 | 86.4 |
| 11 | 314 | 84.3 |
| 12 | 476 | 82.3 |
| **Total** | **7233** | **1119.6** |

**Table S2:** Characteristics of linkage groups within the linkage map constructed based on RADseq data from 160 *T. ivanbureschi* samples from a full-sibling family.

| Primer Pair | Forward Primer Sequence | Reverse Primer Sequence | Product Length (bp) |
|---|---|---|---|
| **CDK-17** | GGCATGGGAAGAACAGAAGA | CCATCTGCTTGGACTGTTGA | 537 |
| **TiY-444315-short** | AGTTCGAGCCAGTACTTTTAGC | CAAACACACGAAAGCACAGTG | 111 |
| **TiY-444315-long** | CACTGTGCTTTCGTGTGTTTG | TGTACTAGAAAGGGTGGGGG | >105 |
| **TiY-137941-short** | GTCACAGCAGCAAATGGTCC | CAGAAGAAGGGCATCTGGG | 104 |
| **TiY-137941-long** | GTCACAGCAGCAAATGGTCC | CCTCTGCTCTGCCTTCACAG | >166 |
| **TiY-95401-short** | CTAGATTCCGGTGAGGCAGG | GGCCCATAGCACCAACATTC | 137 |
| **TiY-95401-long** | GCGTACGGAGTGATTATCCCC | ACACTGCTGCGGAACTGAAG | >199 |
| **TiY-105918-short** | TGAGGATCTGGCTCAATCGC | TCTCCAAAGGTAACGCGCTG | 82 |
| **TiY-105918-long** | AATCTTGTCCACCAGTGTGC | AATTCAGCAGCCCACATGCC | >185 |
| **TiY-442534-short** | AGGGGCATAAGTGGAGGGAC | AGGGTCTGAAAAGGGCCATC | 114 |
| **TiY-416318-short** | TGGGTTTCCAAGTCTCCTCAG | ACTTTCAAGAGTAAGGAGCAGAAG | 89 |
| **TiY-416318-long** | TGGGTTTCCAAGTCTCCTCAG | TGGAGGCCTGAAGTAATAAGCC | >166 |
| **TiY-254147-short** | CCGGTCACATCTCCTTCGAG | GACTGGGCTTGAGAGTCTCG | 150 |
| **TiY-254147-long** | CCGGTCACATCTCCTTCGAG | TCGAAGCAGATGTGACTGGG | >163 |
| **TiY-384959-short** | TGCAGCACAGCAGTAGACTC | CCTTCTCGCATGGACCCTAC | 92 |
| **TiY-384959-long** | GTAGGGTCCATGCGAGAAGG | AGGTGTCGTGTGCCTACTTC | >168 |
| **TiY-201098-short** | TAAACCAGCAAAGCCACCAC | TGTACAATTCCTGCGTAACCG | 83 |
| **TiY-201098-long** | CCACCCCCAAGCACTTAAAG | TGTGTGGGTCCCAAAAGTGG | >200 |
| **TiY-817010-short** | TCTGCTTTGTGTCTGAAGCTTG | TGTGTGTTCCTGTTGGGCTG | 127 |
| **TiY-817010-long** | TCACCTACCACCACAGTTGC | CACTCCTGACTATGGGCCTG | >186 |
| **TiY-301991-short** | GGGGAGTCAGGGTTGTCATG | TCTACTAGCTCACAGGGCAC | 87 |
| **TiY-301991-long** | GGGGAGTCAGGGTTGTCATG | TGGGGTTTCCTACTCAGCTG | >194 |
| **TiY-106308-short** | AGCAAGTTCCAGGAGCTTCC | AGAGCACATGAAGGACCAGC | 125 |
| **TiY-106308-long** | TCACCAGCAGAGTTTCTCCG | TGAAGGACCAGTGGATGCTG | >186 |

**Table S3:** Sequences of all primers used in this study, CDK-17 is an autosomal marker used as a control, all others are candidate Y-linked markers developed for *T. ivanbureschi*.
